# Predicting protein thermal stability changes upon point mutations using statistical potentials: Introducing *HoTMuSiC*

**DOI:** 10.1101/038554

**Authors:** Fabrizio Pucci, Raphael Bourgeas, Marianne Rooman

**Affiliations:** Department of BioModeling, BioInformatics & BioProcesses, Université Libre de Bruxelles, CP 165/61, Roosevelt Ave. 50, 1050 Brussels, Belgium

## Abstract

The accurate prediction of the impact of an amino acid substitution on the thermal stability of a protein is a central issue in protein science, and is of key relevance for the rational optimization of various bioprocesses that use enzymes in unusual conditions. Here we present one of the first computational tools to predict the change in melting temperature Δ*T_m_* upon point mutations, given the protein structure and, when available, the melting temperature *T_m_* of the wild-type protein. The key ingredients of our model structure are standard and temperature-dependent statistical potentials, which are combined with the help of an artificial neural network. The model structure was chosen on the basis of a detailed thermodynamic analysis of the system. The parameters of the model were identified on a set of more than 1,600 mutations with experimentally measured Δ*T_m_*. The performance of our method was tested using a strict 5-fold cross-validation procedure, and was found to be significantly superior to that of competing methods. We obtained a root mean square deviation between predicted and experimental Δ*T_m_* values of 4.2°C that reduces to 2.9°C when ten percent outliers are removed. A webserver-based tool is freely available for non-commercial use at *soft.dezyme.com*.

## Introduction

The possibility of rationally modifying protein sequences to increase their thermal resistance is a main goal in protein engineering. Indeed, the design of new enzymes and other proteins that remain stable and active at temperatures that differ from their physiological temperatures would allow the optimization of a wide series of biotechnological processes in many sectors such as agro-food, biopharmaceuticals and environment.

Unfortunately, it is extremely complicated to predict the effect of mutations on the thermal stability of proteins, defined through the melting temperature *T_m_*, *i.e.* the temperature at which the protein undergoes the reversible (un)folding transition. It is even more difficult than predicting the change in thermodynamic stability defined by the folding free energy Δ*G(T_r_)* at room temperature (*T_r_*), since it requires a precise understanding of the variation of the free energy Δ*G*(*T*) as a function of the temperature (*T*) of the different types of interactions that contribute to protein stability, *i.e.* between the various chemical groups that form the solvent and the 20 amino acids. This is a longstanding problem that is currently far from being solved. The analyses performed in the last thirty years led to the conclusion that there is no unique and specific factor that ensures an enhancement of the thermal stability of all proteins, but that there is - though very approximately - such a factor inside each protein family, as homologous proteins tend to involve the same kinds of stabilizing interactions (see for example [1–10] and references therein).

A series of experimental approaches have been developed to design new mutants with higher or lower melting temperature than the wild-type protein. They are mostly based on directed evolution experiments that mimic natural evolution, sometimes in combination with computational approaches (see [11, 12] and references therein). Unfortunately, these methods are only partially successful. Indeed, they are expensive and time-consuming, and moreover limited by the vastness of sequence space and the low frequency of the thermally stabilizing mutations.

In *silico* protein engineering constitutes an alternative for the design of new proteins with modified stability. Different computational tools based on a variety of approaches and information including residue conservation, energy estimations as well as structural, sequence and dynamical features, have been developed to get a prediction of the thermodynamic stability changes upon point mutations defined through ΔΔ*G(T_r_)*, the difference of folding free energy Δ*G* between the mutant and wild-type proteins at room temperature. Quite interestingly, some of these methods can reach a good accuracy at very low computational cost, with the sole knowledge of the wild-type structure [13–23] (see also [24] for a comparison of their performances). This makes the fastest among them ideal tools for stability analyses of the entire structurome. One of the major drawbacks of these methods is that the results are usually biased towards the training datasets even if strict cross validation is applied, which makes the estimation of their true performances an almost impossible task [25].

The impact of point mutations on the thermal stability, defined through Δ*T_m_* which measures how the protein melting temperature changes upon mutations, has been much less investigated than the thermodynamic stability, as it is more intricate and requires taking into account that the amino acid interactions are temperature dependent. Therefore, there are very few *in silico* tools for predicting Δ*T_m_* [16, 26–28]. The common strategy to study the enhancement of thermal resistance is to assume the thermodynamics and thermal stabilities to be perfectly correlated (or ΔΔ*G(Tr)* and Δ*T_m_* to be perfectly anticorrelated). Unfortunately, even if this approximation can be used in a first instance, it is not always reliable [29]. As a consequence, the predictions of Δ*T_m_* are in general less accurate because the intrinsic errors on the ΔΔ*G* predictions have to be summed with the errors due to the imperfect correlation of the two stabilities. As an example, the anticorrelation between ΔΔ*G*’s predicted using the thermodynamic stability change predictor PoPMuSiC [22] and measured Δ*T_m_*’s is not so satisfactory and is equal to 0.51, whereas the correlation between predicted and experimental ΔΔ*G*’s is 0.63.

For all these reasons, it is necessary to design a specific computational tool for predicting Δ*T_m_* in a fast and more precise way. This is the aim of the present paper, in which we present a new, knowledge-and thermodynamics-based, method called HoTMuSiC, which is able to predict this quantity using as sole input data, the three-dimensional (3D) structure of the wild-type protein and – when available – its melting temperature *T_m_*. A very preliminary version of this method has been published in [30]. The main reasons of the success of HoTMuSiC are rooted on the one hand in the thorough physical analysis of the system which helped correct guessing the form of the model structures, and on the other hand in the use of temperature-dependent statistical potentials [22, 31–33] that are extracted from non-redundant datasets of protein X-ray structures of thermostable and mesostable proteins. We would like to emphasize that these are presently the only potentials that yield an estimation of the temperature dependence of the folding free energy contributions of the different amino acid interactions - albeit in an approximate, effective, manner.

## Results

### Theoretical analysis

The thermodynamic stability change upon mutation is measured by ΔΔ*G(T_r_), i.e.* the difference between the Gibbs folding free energies of the mutant (Δ*G^mut^*) and wild-type (Δ*G^Wild^*) proteins at the reference temperature *T_r_*:

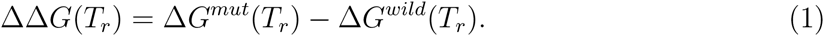

Usually *T_r_* is taken as the room temperature: *T_r_ = 298K. ΔΔG*’s upon point mutations can be predicted in *silico* using a series of tools [13–23], which reach a relatively good accuracy with a standard deviation between the experimental and predicted ΔΔ*G*’s of 1 to 2 kcal/mol.

Less prediction methods have been developed for the change in thermal stability upon mutations, measured by Δ*T_m_, i.e.* the difference between the melting temperature of the mutant 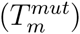 and wild-type 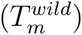 proteins:

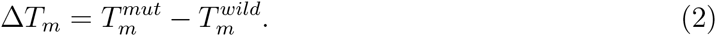

In a first approximation the two protein stabilities can be assumed to be strongly interdependent. Indeed, focusing on two-state folding transitions and assuming: (1) the mutations to be small perturbations with respect to the wild-type state; (2) the folding heat capacity Δ*C_P_* to be *T*-independent; (3) its variations upon mutations to vanish 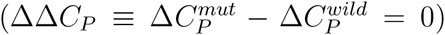 and (4) similarly for the folding enthalpy 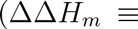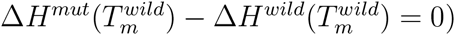, one can derive the simple relation:

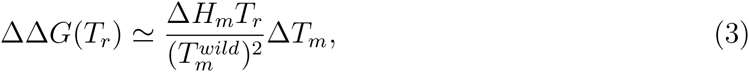

with 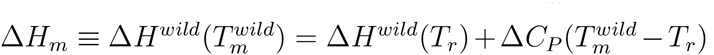. This equation directly relates the two stabilities, however with a coefficient that depends on the thermal characteristics of the wild-type protein through its melting temperature 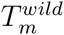 and its folding free enthalpy Δ*H_m_.* Note that, since the enthalpy change upon folding is negative, ΔΔ*G* and Δ*T_m_* are anticorrelated, as expected.

Unfortunately, the situation is in general less simple, especially for highly destabilizing or highly stabilizing mutations, and we have to take into account that the enthalpy and heat capacity variations do not vanish, *i.e. ΔΔH_m_ ≠ 0 ≠ ΔΔCP*. This is illustrated by the fact that the correlation coefficient between experimental Δ*T_m_*’s and ΔΔ*G*’s is about —0.8, which signals an imperfect correlation between these quantities. In this case, and still assuming Δ*C_P_* to be *T*-independent, Eq. (3) becomes:

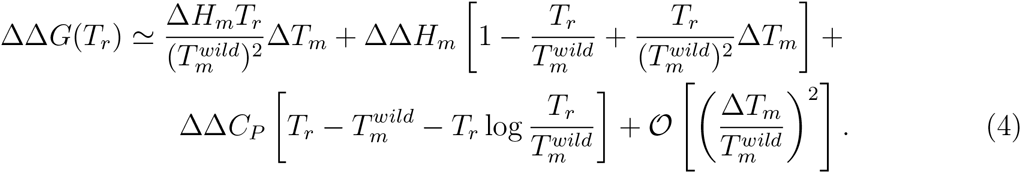

Thus the simple relation between the two descriptors of protein stability is lost: the proportionality coefficient can be positive or negative, and becomes also mutation-dependent in addition of being protein-dependent.

It is easier to understand the meaning of the possible types of correlations between the two stabilities with a graphical representation. If the assumption of small perturbation and as a consequence Eq. (3) holds, the full protein stability curve changes upon mutation like in Fig. 1.a. Typically, in this case, the wild-type has one interaction more than the mutant (or conversely). Otherwise the scenario is more similar to Fig. 1.b, where one cannot say *a priori* which type of connection there is between ΔΔ*G(T_r_)* and Δ*T_m_* without the knowledge of additional information. It can for example occur when an interaction that is more stabilizing at room temperature is replaced by another that is more *T*-resistant thereby modifying the Δ*H_m_,* or when a change in the 3D structure occurs which modifies the protein’s solvent accessible surface area and thus the Δ*C_P_*.

**FIG. 1:**
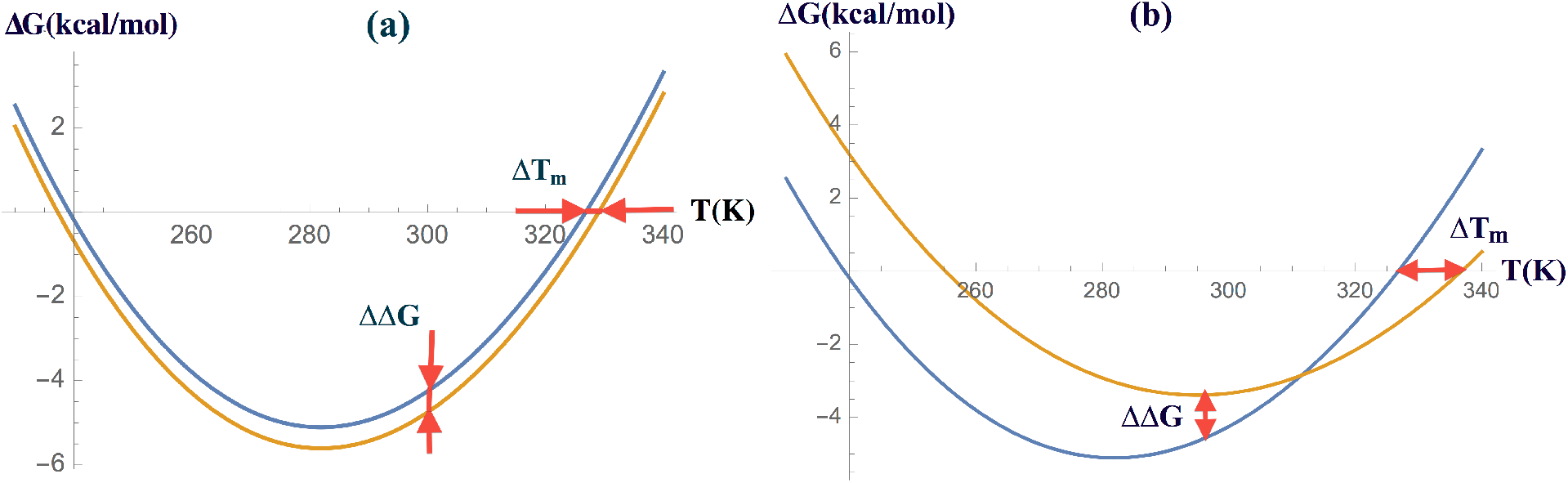
Effect of mutations on the full stability curve of an hypothetical protein. (a) For a small perturbation of the wild-type state, we observe a negative proportionality between Δ*T_m_* and ΔΔ*G*; (b) Such simple relation is lost for highly destabilizing or stabilizing mutations.

In a first approach to the prediction of Δ*T_m_* upon mutations, we consider the small perturbation approximation and thus Eq. (3) as valid and compute Δ*T_m_* as the sum of usual, temperature-independent, statistical potential contributions. In a second approach, which is expected to be more accurate for highly destabilizing or stabilizing mutations, we use ΔΔ*G*-values calculated at different temperatures. Note however that structural modifications are more likely to occur in such case, and thus that the accuracy of the energy evaluations on the basis of the wild-type structure alone is questionable. For the purpose of estimating ΔΔ*G(T)*, we use the formalism of the temperature-dependent statistical potentials introduced in [31–33].

### Construction of the model

#### Standard and temperature-dependent statistical potentials

The standard formalism of statistical potentials [34–36] has been fruitfully applied to a variety of analyses that range from the prediction of protein structure, stability, and proteinprotein and protein-ligand binding affinities. It basically consists in deriving a potential of mean force (PMF) from frequencies of associations of structure and sequence elements in a dataset of protein X-ray structures. Under some approximations whose validity has been discussed [36–40] and making use of the Boltzmann law, the simplest PMF can be written as:

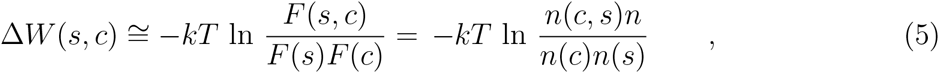

where *c* and *s* are structure and sequence elements respectively, *F* represent the relative frequencies of *c* and/or *s* which are expressed as a function of the number of occurrences *n*, *k* is the Boltzmann constant and *T* the absolute temperature. Following [41], higher order potentials can be constructed by considering more than two structure elements and/or sequence elements. Considering for example two sequence elements *s* and *s’* and one structure element *c*, the above expression of PMF gets modified as:

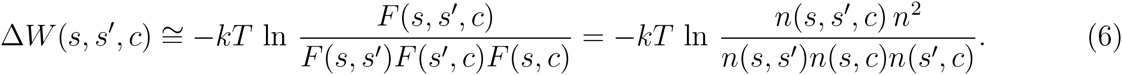

Using Eqs (5-6) and their generalizations defined in [41], we derived 9 different statistical potentials from a dataset of about 4,100 proteins with well-resolved 3D structure and low sequence similarity; they are listed in Table I. They differ by the number of sequence and/or structure elements involved and by their type. Each sequence element *s* is an amino acid type at a given position and each structure element *c* is either the spatial distance *d* between two residues, the backbone torsion angle domain *t* or the solvent accessibility *a* of a residue.

**TABLE I:** List of 9 *T*-independent and 5 *T*-dependent statistical potentials used in the Δ*T_m_-*prediction methods. The subscripts *M/T* indicate that the potentials are extracted from either the mesostable (*M*) or thermostable (*T*) protein subset.

| Type | Standard potentials | $T$-dependent potentials |
| --- | --- | --- |
| Distance | $\Delta W_{[sd]} ; \Delta W_{[sds]}$ | $\Delta W_{[sd]}^{M/T} ; \Delta W_{[sds]}^{M/T}$ |
| Distance/Accessibility | $\Delta W_{[sad]}$ | $\Delta W_{[sad]}^{M/T}$ |
| Distance/Torsion | $\Delta W_{[std]}$ | - |
| Accessibility | $\Delta W_{[sa]} ; \Delta W_{[saa]}$ | $\Delta W_{[sa]}^{M/T} + \frac{1}{2}(\Delta W_{[saa]}^{M/T} + \Delta W_{[ssa]}^{M/T})$ |
| Torsion | $\Delta W_{[st]} ; \Delta W_{[stt]} ; \Delta W_{[sst]}$ | $\Delta W_{[st]}^{M/T} + \frac{1}{2}(\Delta W_{[stt]}^{M/T} + \Delta W_{[sst]}^{M/T})$ |

The statistical potential formalism has recently been extended to include more properly the thermal characteristics of proteins and in particular the fact that amino acid interactions are temperature-dependent [9, 31–33]. Following this approach, a dataset of about 170 proteins with known melting temperature and 3D structure was used and divided into two subsets, one with only mesostable proteins (with *T_m_* less than about 65°C), and the other with thermostable proteins (with *T_m_* higher than about 65°C). A series of 9 different potentials were extracted from each subset, which are listed in Table I. To limit the number of parameters to be optimized (see next section) and thus to avoid overfitting as much as possible, these twelve potentials were combined into five potentials (Table I).

As expected, these *T*-dependent potentials reflect the thermal characteristics – mesostable or thermostable – of the subset from which they are derived: the former set better describes the interactions in the low temperature region while the second set better reflects the thermal properties at high temperatures. This approach has shown good performances in the prediction of the thermal resistance and of the stability curve as a function of the temperature of proteins that belong to the same homologous family [32, 33].

The structure elements defining these potentials are the same as for the temperature-independent potentials. In contrast, the size of the mesostable and thermostable protein datasets is much smaller than the dataset used for the standard statistical potentials, as it requires the knowledge of the melting temperature. To deal with the smallness of the datasets, we used several tricks that consists of corrections for sparse data and the smoothing of the potentials, following [9, 31–33].

In addition to the standard and *T*-dependent potentials Δ*W,* we also considered volume terms in the folding free energy estimation, which are defined as the volume difference Δ*V* between the mutant and wild-type amino acids [22, 23]. In order to take into account that the creation of a cavity in the protein interior (Δ*V* < 0) can have a different impact on the stability compared to the addition of stress (Δ*V* > 0), we introduced two separate terms (Δ*V_−_*) and (Δ*V*_+_) defined as 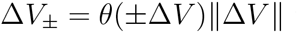 where θ is the Heaviside function.

#### Artificial neural networks and parameter identification

The above-defined potential terms were combined to predict how the melting temperature changes upon mutations, using two different model structures. The second model (called *T_m_*-HoTMuSiC) uses the *T_m_* of the wild-type, while the first (HoTMuSiC) does not. To identify the parameters involved in these combinations, we used an artificial neural network (ANN) with peculiar activation functions.

In the first approach (HoTMuSiC), we assumed that Δ*T_m_* can be written as the sum of twelve contributions, the nine energy terms 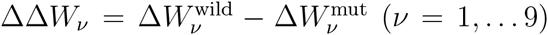 computed from the standard, *T*-independent, statistical potentials listed in Table 1, the two volume terms and an “independent” term that only depends on the solvent accessibility. The functional form reads as:

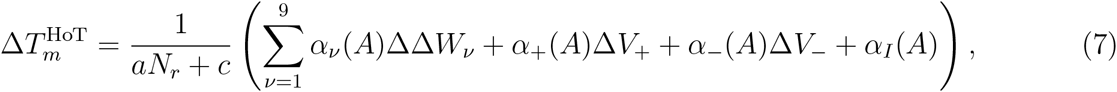

where N_r_ is the number of residues in the protein, and the coefficients *α_v_*(A) were chosen to be sigmoid functions of the solvent accessibility A of the mutated residues:

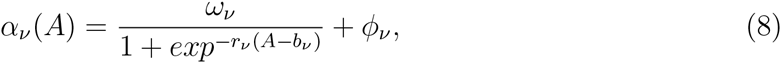

with 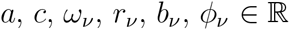. We have chosen the activation functions to be sigmoidal since they model a smooth transition from the protein core to the surface, and since the weight of the different energy contributions have been shown to differ in these two regions [22][47].

To identify the fifty parameters introduced in Eq. (7), we have chosen a standard feed-forward ANN with one input and one output layer (schematically depicted in Fig. 2a). The cost function to be minimized is the mean square deviation between the experimental and predicted values of Δ*T_m_* for the dataset *S_mut_* that contains *N*_mut_=1,626 mutations for which an experimental Δ*T_m_-*value is available (see Methods section):

**FIG. 2:**
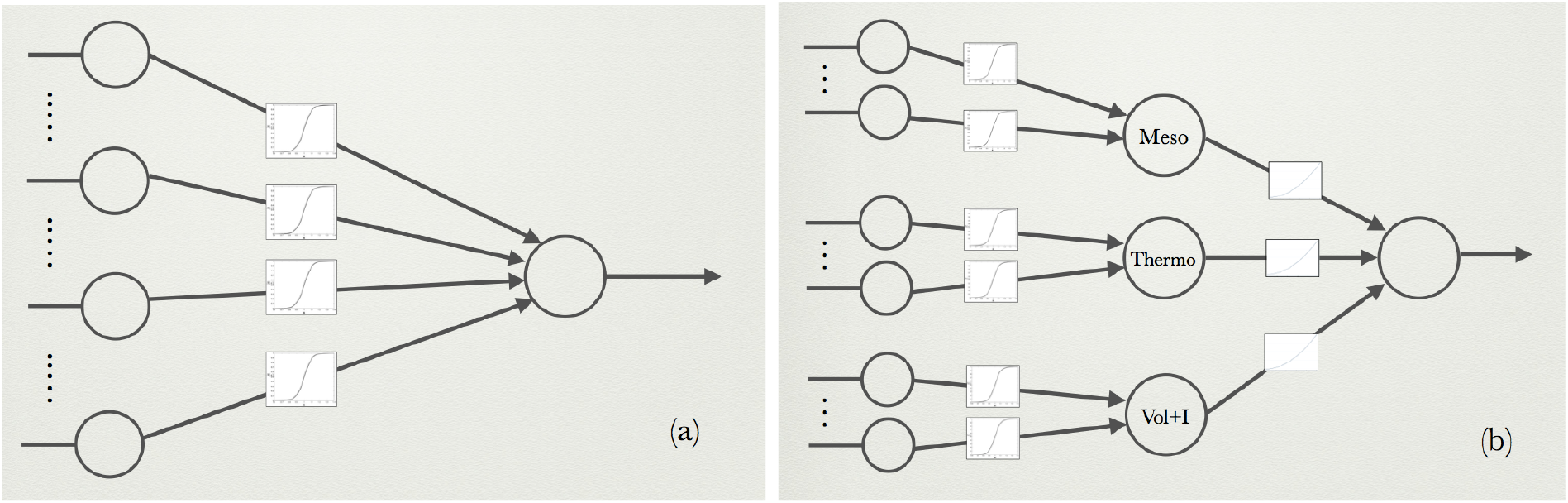
Schematic representation of the ANN’s used for the parameter identifications. (a) HoTMuSiC method: 2-layer ANN, corresponding to a perceptron with sigmoid activation functions and 12 input neurons encoding the 9 *T*-independent potentials specified in Table I, two volume terms and an independent term; (b) *T*_m_-HoTMuSiC method: 3-layer ANN, consisting of 3 perceptrons with sigmoid weights; the first perceptron has 5 input neurons encoding the 5 mesostable potentials listed in Table I, the second has 5 input neurons corresponding to the 5 thermostable potentials, and the last perceptron has 3 neurons for the volume and independent terms. The outputs of these three perceptrons (Meso, Thermo, and Vol+I) are the inputs of another perceptron with polynomial weight functions.

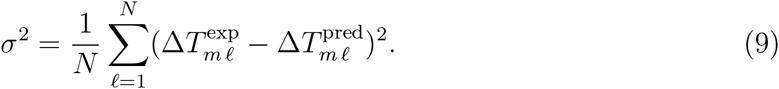

The initial values of the weights were chosen randomly. To take into account that the ANN training algorithm can get stuck in local minima, the initialization and training processes were repeated about thirty times and the solution reaching the lowest *σ*-value was chosen [43, 44].

We used a strict five-fold cross validation procedure with an early stopping procedure for evaluating the method’s performance. More precisely, the entire set of mutations was split randomly into a training set (containing 90% of the mutations) and a test set (with the remaining10%). The training set was then further randomly split into a smaller set (with 80% of the mutations) on which the parameters were identified, and a validation set (with 10% mutations) on which the early stop threshold was determined, namely the maximum number of iterations in the gradient descent procedure before its convergence [45, 46]. This early stop is necessary to avoid overfitting, as in general the network starts to learn too much from the training dataset after a certain number of iterations, with the consequence that the error in cross validation starts to raise. As a final step in the computation, the prediction error *σ* is independently calculated on the test set.

In the second method for predicting Δ*T_m_*, called *T_m_-*HoTMuSiC, we used quite a different approach, with as building blocks the *T*-dependent statistical potentials listed in Table I and the 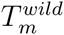 of the wild-type protein. More precisely, Eq. (7) describing the Δ*T_m_* functional was modiied into:

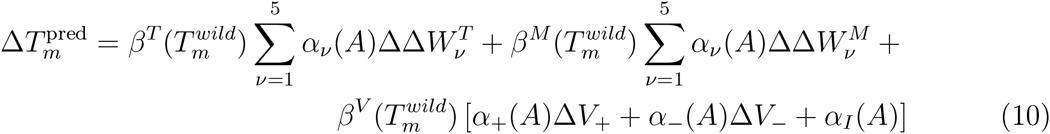

where the *α_v_ (A)* parameters are sigmoid functions of the solvent accessibility A (Eq. (8)) and the three functions 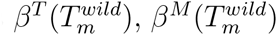 and 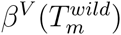 are polynomial functions of the melting temperature of the wild-type protein and its number of residues *N_r_.* Their functional form is guessed from Eq. (3) and approximated as:

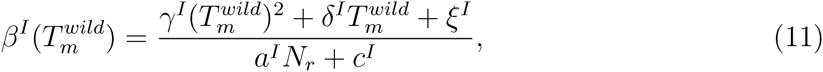

with *I=T* (thermostable), *M* (mesostable) or *V* (volume). The dependence on the number of residues *N_r_* comes from the enthalpy factor Δ*H_m_* in Eq. (3), as these two quantities show a good correlation in a first approximation [48].

To identify the 67 parameters of this second method, an ANN with an input layer, a hidden layer and an output layer is used, as shown schematically in Fig. 2b. The input layer consists of three sets of neurons, one set encoding the mesostable potentials, a second one the thermostable potentials, and a third one the volume and independent terms. Three perceptrons use these three sets of input neurons and generate the three output signals of the hidden layer. These are the input of yet another perceptron, which yields the Δ*T_m_-* prediction as output. The initialization and identification procedures of all the weights and the cross validation procedure are the same as for the first method.

The final Δ*T_m_* predictions of the *T_m_-*HoTMuSiC method are defined as the mean of the two predictions given by Eqs (10) and (7):

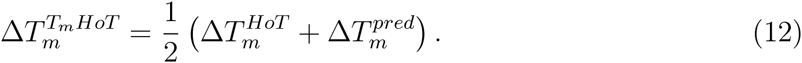

### Performance of HoTMuSiC and *T_m_-*HoTMuSiC

The values of the root mean square deviation *σ* between measured and predicted Δ*T_m_* values (Eq. 9), computed in strict cross validation, are shown in Table II. For HoTMuSiC, we obtained *σ =* 4.3°C; the Pearson correlation coefficient *r* between experimental and predicted Δ*T_m_*’s is 0.59. The performance of the *T_m_-*HoTMuSiC version is slightly better with Δ=4.2°C and *r* = 0.61. When ten percent outliers are excluded, *σ* decreases to 2.9°C and *r* rises to 0.75. The results are plotted in Fig. 3.

**TABLE II:**
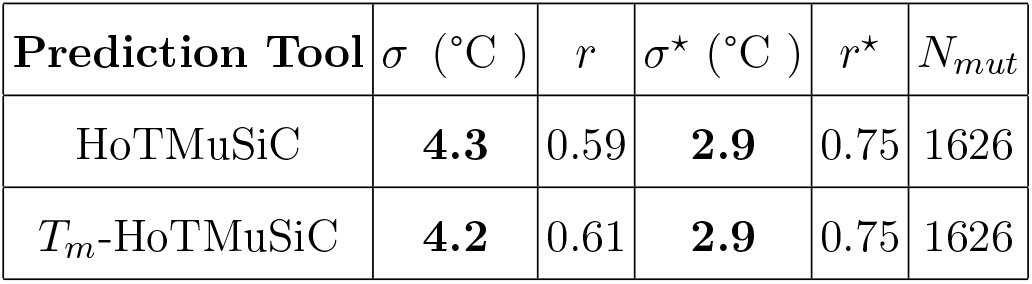
Scores of HoTMuSiC and *T_m_-*HoTMuSiC; *σ*\* and *r*\* correspond to *σ* and *r* with 10% outliers removed.

| Prediction Tool | $\sigma$ ($^{\circ}\text{C}$) | $r$ | $\sigma^*$ ($^{\circ}\text{C}$) | $r^*$ | $N_{mut}$ |
| --- | --- | --- | --- | --- | --- |
| HoTMuSiC | **4.3** | 0.59 | **2.9** | 0.75 | 1626 |
| $T_m$-HoTMuSiC | **4.2** | 0.61 | **2.9** | 0.75 | 1626 |

**FIG. 3:**
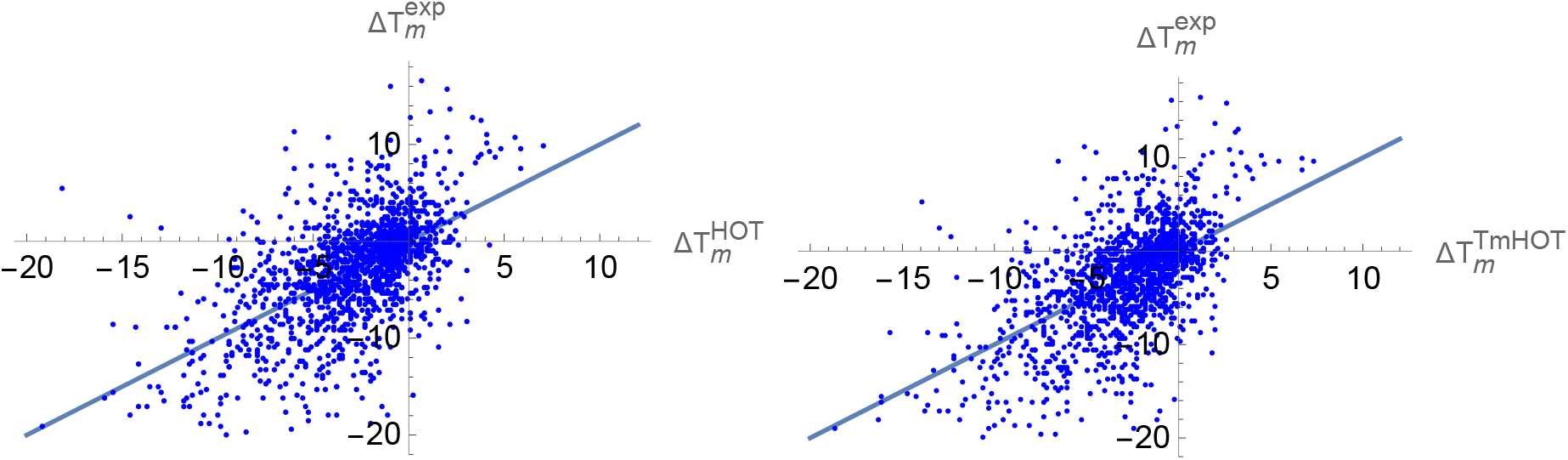
Experimental Δ*T_m_* values versus predicted 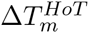 (left, *r*=0.59) and 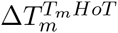 (right, *r* = 0.61) values. The straight lines are the bisectors of the first and third quadrants. The temperatures are in Celsius degrees (°C).

The *T_m_-*HoTMuSiC version that encodes information about the melting temperature of the wild-type protein and uses *T*-dependent potentials thus performs slightly better than the *T*-independent version, but not as much as could be expected on the basis of earlier analyses [32][33]. Indeed, in principle, the mesostable potentials should describe the mutations in the mesostable proteins much better than the thermostable potentials and vice-versa. The reasons for this mitigated result could be due to the lower accuracy of the *T*-dependent potentials compared to the standard ones since they are extracted from smaller protein sets. Or they could be rooted in the information loss upon the reduction of the number of potentials (see Eq. (7) versus Eq. (10)), which is done to avoid overfitting issues.

Moreover, we can see from Table III and Fig. 4 that the *σ*-value for amino acid substitutions in the core (A<15%) and partially buried positions (15%<A<50%) is on the average larger than that of surface mutations (A>50%). In contrast, the correlation coefficient r is higher in the core and in the partially buried region and smaller at the surface. This apparent discrepancy is in fact due to the higher variance of Δ*T_m_* in the core, which drives the correlation and increases the value of *r*. In the surface region, the predictions are more accurate (lower *σ*) but the Δ*T_m_* variance and the correlation coefficient are lower. When normalizing *σ* by the standard deviation of Δ*T_m_*, we obtain values that increase from the core to the surface (see Table III).

**TABLE III:** Scores of *T_m_*-HoTMuSiC as a function of the solvent accessibility A of the mutated residues.

| **Solvent accessibility** | $\sigma$ (°C) | $r$ | $\frac{\sigma}{\sqrt{\langle(\Delta T_m)^2\rangle}}$ | $N_{mut}$ |
| --- | --- | --- | --- | --- |
| A<15% | 4.9 | 0.70 | 0.66 | 734 |
| 15%<A<50% | 4.2 | 0.57 | 0.81 | 513 |
| A>50% | 2.8 | 0.54 | 0.83 | 379 |

**FIG. 4:**
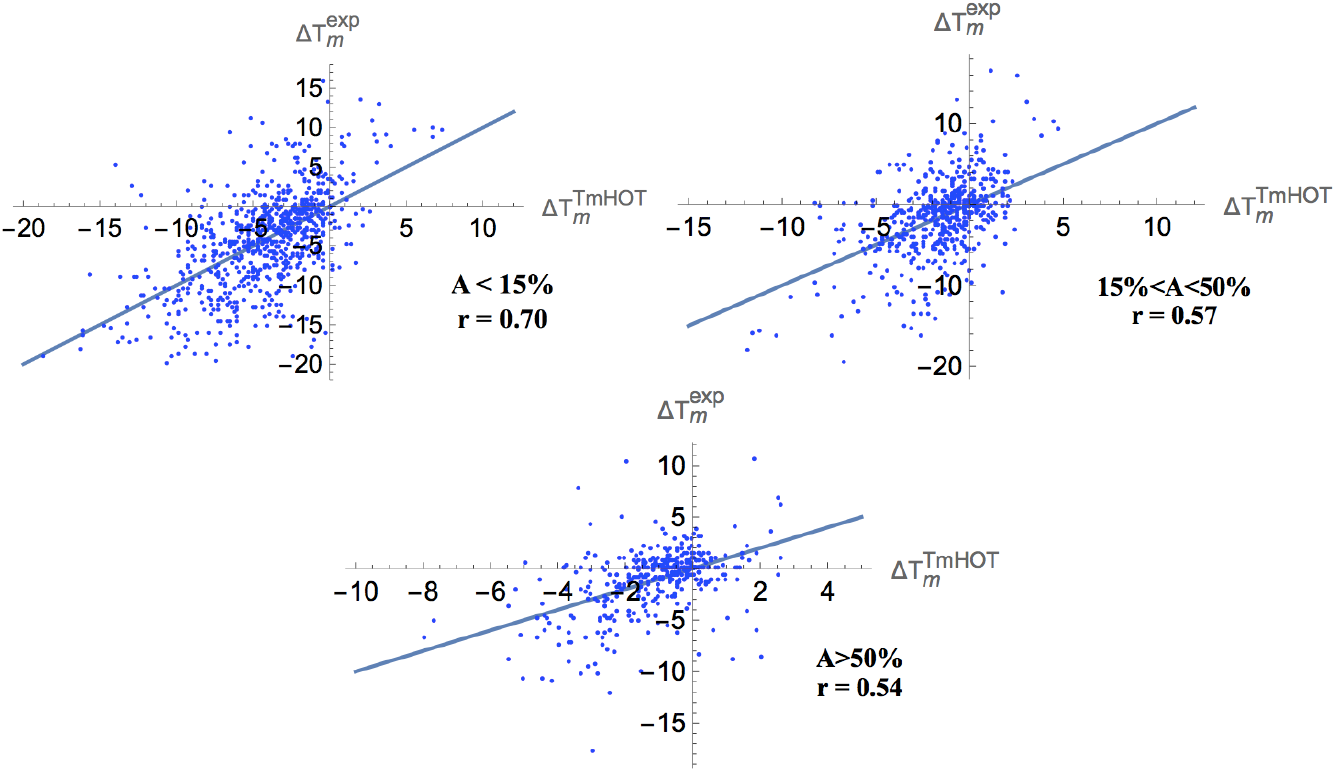
Experimental Δ*T_m_*’s versus predicted 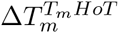 values for mutations in different protein regions: core (*A* <15%), partially buried (15%<A<50%), and surface (*A* >50%). The straight lines are the bisectors of the first and third quadrants. The temperatures are in °C.

### Comparison with other Δ*T_m_* predictors

As far as we known, only two other Δ*T_m_* predictors are described in the literature, which are strongly different from ours. The strategy of Saraboji et al. [28] consists in predicting for a mutation from wild-type residue W to mutant M the mean value of the Δ*T_m_*’s of all the analogous mutations W→M in the training dataset. A similar strategy proposed in the same work is based on the classification of the mutations in terms of the secondary structure and solvent accessibility of the wild-type residues and predicts as Δ*T_m_* the mean of the experimental Δ*T_m_*’s occurring in the suitable class in the learning set. A limitation of this approach is that not all wild-type to mutant mutations are present in the learning set due to the lack of experimental data. The second method is called AutoMute [16, 26, 27] and is based on residue environment scores. It proceeds by reducing protein 3D structures to ensembles of *C_α_* atoms, and applying Delaunay tessellation to identify quadruplets of nearest neighbor residues. A log-likelihood potential is constructed from the number of occurrences of the quadruplets in a dataset of 3D structures and then used as the key ingredient in the computation of Δ*T_m_*.

To make a cross-validated comparison between our results and those of these two methods, we have chosen a subset *S_sub_* of our dataset *S_mut_* consisting of the mutations that are not present in the AutoMute learning set (see [60] for a list), and trained versions of *T_m_*-HoTMuSiC and the Saraboji method on the set *S_mut_ \ S_sub_*. The performances of the three methods are reported in Table IV. *T_m_-*HoTMuSiC shows the best performance, with an improvement of about 20% and 30% with respect to AutoMute [16] and the Saraboji method [28], respectively. The σ-values computed on the *S_sub_* set are equal to 3.7, 4.7 and 5.4 °C for *T_m_-*HoTMuSiC, AutoMute and Saraboji method, respectively. Note that the performance of the latter two methods has been evaluated on a slightly reduced dataset, since the Δ*T_m_* values could not be computed for some of the mutations.

**TABLE IV:** Comparison between the performances of HotMuSiC and those of the two other methods evaluated in cross-validation on the dataset *S_sub_*.

| **Prediction Tool** | $\sigma$ (°C) | $r$ | $N_{mut}$ |
| --- | --- | --- | --- |
| $T_m$-HoTMuSiC | 3.7 | 0.65 | 630 |
| AutoMute v2.0 [16] | 4.7 | 0.42 | 607 |
| Saraboji [28] | 5.4 | 0.25 | 580 |

## Discussion

We developed a thermodynamics-based and knowledge-driven Δ*T_m_* prediction method that does not exploit the common assumption of a perfect correlation between thermal and thermodynamics stabilities. The basic ingredients of our approach include standard and *T*-dependent statistical potentials that are combined through the use of ANN s. The performance in cross validation of the two versions of our method, HoTMuSiC and *T_m_-*HotMuSiC (which requires the *T_m_* of the wild type), are quite good with a *σ*-value of 4.3 and 4.2 °C, respectively, which goes down to 2.9 °C upon removal of 10% outliers. They perform significantly better, by 20 to 30%, than the two other Δ*T_m_* prediction methods described in the literature.

HoTMuSiC and *T_m_*-HotMuSiC are accessible via the webserver soft.dezyme.com and are free for non-commercial use. They are extremely fast and allow the Δ*T_m_* predictions for all possible single-site mutations in a protein in a few minutes. This webserver is presented in an application note [49].

Our software thus yields quite accurate results, and allows rapid screening of all possible point mutations in a protein structure and identifying a subset that is likely to yield the required thermal resistance. This subset must then be analyzed further, either by using more detailed computational techniques, or by experimental means. HoTMuSiC and *T_m_-*HotMuSiC are very useful and user-friendly tools for every researcher who wishes to rationally design modified proteins with controlled characteristics.

Notwithstanding the large applicability and good accuracy of HoTMuSiC and *T_m_-*HotMuSiC, it is worth discussing their limitations and the sources of errors that affect the predictions. These are:

- The wild-type and mutant structures are supposed to be identical (up to the substituted side chain) and the possible structural modifications are only encoded in the volume terms Δ*V_±_*; local structure rearrangements upon residue substitutions, for example in the hydrophobic core, yet depend on many more parameters such as the residue depth and the backbone flexibility [50–54].
- The mutation dataset is strongly unbalanced towards destabilizing mutations, which is likely to add unwanted hidden biases even if a strict cross validation procedure is applied [25]. Only when more stabilizing mutations will be experimentally characterized will we be able to completely exclude the biasing impact of this stabilizing-destabilizing asymmetry.
- The experimental conditions at which the *T_m_* measurements are performed usually differ in terms of pH, ion concentration or buffer composition, which induces noise in the learning set and errors in the predictions. To limit this problem, we chose the entries derived from experiments performed at pH as close as possible to seven, and made a weighted average of the experimental Δ*T_m_*’s of a same mutation, when available.
- The *T*-dependent potentials suffer from the smallness of the dataset of protein structures with known *T_m_*. Different tricks have been used to limit this issue.
- The possible parameter overfitting is like always an important concern, especially for the *T_m_-*HoTMuSiC version, in which the number of potentials is three times larger than for HoTMuSiC. To avoid overfitting, we decided to decrease the number of parameters in *T_m_-*HoTMuSiC by fixing some coefficients of the linear combination of potentials (see Table I).

Different ways will be explored in an attempt to further improve the prediction performances of HoTMuSiC. They obviously include the enlargement of the datasets of proteins of known structure and *T_m_*, and of the mutations of known Δ*T_m_*. We will also investigate different ANN architectures, the addition of hidden layers, and the inclusion of other features such as the change in conformational flexibility upon mutation, which seems related to the thermal stability even if a quantitative connection between the two quantities on a large scale is still missing [55–58]. Finally, it could be worth analyzing the wrong predictions in view of identifying the factors that should be taken into account to make HoTMuSiC even more performing.

## Methods

### Set of experimentally characterized mutations

We started collecting the mutations with experimentally measured Δ*T_m_* value from the ProTherm database [59] and the literature. Each entry was then manually checked from the original literature to remove errors and select those that satisfy the following criteria: were only considered (1) mutations in monomeric proteins of known X-ray structure with resolution below 2.5 Å, (2) mutations whose experimental Δ*T_m_* was measured in absence of chemical denaturants, (3) simple two-state (un)folding transitions, and (4) single point mutations. Destabilizing or stabilizing mutations by more than 20 °C were overlooked, as they probably induce important structural modifications that our method is unable to model. Using this procedure, we obtained a set *S_mut_* of 1,626 mutations that belong to 90 proteins and have an experimental Δ*T_m_*. More information and their list can be found in [60].

## Acknowledgements

FP is Postdoctoral researcher, RB Postdoctoral Fellow and MR Research Director at the Belgian Fund for Scientific Research (FNRS). Financial support from the FNRS through an FRFC grant is acknowledged.

## Author contributions statement

F.P. and M.R. conceived the experiment(s), F.P. and R.B. conducted the experiment(s), F.P. and M.R. analyzed the results, F.P. and M.R. wrote the paper. All authors reviewed the manuscript.

## Additional information

The authors declare no competing financial interests.

## References

[1] Vogt G, Woell S, Argos P, Protein thermal stability, hydrogen bonds, and ion pairs, J. Mol. Biol. 269, 631–43 (1997).

[2] Kumar S, Tsai CJ, Nussinov R, Thermodynamic differences among homologous thermophilic and mesophilic proteins, Biochemistry 40, 14152–65 (2001).

[3] Kumar S, Tsai CJ, Nussinov R, Factors enhancing protein thermostability, Protein Eng. 13, 179–91 (2000).

[4] Kumar S, Nussinov R, Salt bridge stability in monomeric proteins, J. Mol. Biol. 293, 1241–55 (1999).

[5] Kumar S, Nussinov R, Close-range electrostatic interactions in proteins, ChemBioChem. 3, 604–17 (2002).

[6] Thompson MJ, Eisenberg D, Transproteomic evidence of a loop- deletion mechanism for enhancing protein thermostability, J. Mol. Biol. 290, 595–604 (1999).

[7] Chakravarty S, Varadarajan R, Elucidation of factors responsible for enhanced thermal stability of proteins: a structural genomics based study, Biochemistry 41, 8152–61 (2002).

[8] Berezovsky IN, The diversity of physical forces and mechanisms in intermolecular interactions, Phys. Biol. 8, 035002 (2001).

[9] Folch B, Dehouck Y, Rooman M, Thermo-and mesostabilizing protein interactions identified by temperature-dependent statistical potentials, Biophys. J. 98, 667–77 (2010).

[10] Van Dijk E, Hoogeveen A, Abeln S, The Hydrophobic Temperature Dependence of Amino Acids Directly Calculated from Protein Structures, PLoS Comput Biol 11, e1004277 (2015).

[11] Eijsink VG, Gaeseidnes S, Borchert TV, van den Burg B, Directed evolution of enzyme stability, Biomol Eng 22, 21–30 (2005).

[12] Counago R, Chen S, Shamoo Y, In vivo molecular evolution reveals biophysical origins of organismal fitness, Mol Cell. 22, 441–9 (2006).

[13] Guerois R, Nielsen JE and Serrano L, Predicting changes in the stability of proteins and protein complexes: a study of more than 1000 mutations, J. Mol. Biol. 320, 369–387 (2002).

[14] Parthiban V, Gromiha MM, Schomburg D, CUPSAT: prediction of protein stability upon point mutations, Nucleic Acids Res. 34, W239–W242 (2002).

[15] Seeliger D, De Groot DL, Protein thermostability calculations using alchemical free energy simulations, Biophys. J. 89, 2309–16 (2010).

[16] Masso M and Vaisman II, Accurate prediction of stability changes in protein mutants by combining machine learning with structure based computational mutagenesis, Bioinformatics 24, 2002–2009 (2008).

[17] Capriotti E, Fariselli P, Casadio R, I-Mutant2.0: predicting stability changes upon mutation from the protein sequence or structure, Nucleic Acids Res. 33, W306–W310 (2005).

[18] Huang LT, Gromiha MM, Ho SY, Sequence analysis and rule development of predicting protein stability change upon mutation using decision tree model, J. Mol. Model. 13, 879–890 (2007).

[19] Cheng J, Randall A, Baldi P, Prediction of protein stability changes for single-site mutations using support vector machines, Proteins 62, 1125–1132 (2006).

[20] Potapov C, Cohen M, Schreiber G, Assessing computational methods for predicting protein stability change upon mutation using tree model, J. Mol. Model 13, 879–890 (2007).

[21] Ozen A, Gonen M, Alpaydan E, Haliloglu T, Machine learning integration for predicting the effect of single amino acid substitutions on protein stability, BMC Struct. Biol. 9, 66 (2009).

[22] Dehouck Y, Grosfils A, Folch B, Gilis D, Bogaerts P, Rooman M, Fast and accurate predictions of protein stability changes upon mutations using statistical potentials and neural networks: PoPMuSiC-2.0, Bioinformatics 25, 2537–43 (2009).

[23] Dehouck Y, Kwasigroch JM, Gilis D, Rooman M, PoPMuSiC 2.1: a web server for the estimation of protein stability changes upon mutation and sequence optimality, BMC Bioinformatic 12, 151 (2011),.

[24] Kahn S, Vihinen M, Performance of protein stability predictors, Human Mutation 31, 675–684 (2010).

[25] Pucci, F, Bernaerts K, Teheux F, Gilis D, Rooman, M, Symmetry Principles in Optimization Problems: An Application to Protein Stability Prediction, IFAC Proceedings, MathMod 2015 8, 458–463.

[26] Masso M, Vaisman II, AUTO-MUTE 2.0: A Portable Framework with Enhanced Capabilities for Predicting Protein Functional Consequences upon Mutation, Advances in Bioinformatics, ID 278385 (2014).

[27] M Masso, I.I Vaisman AUTO-MUTE: Web-based tools for predicting stability changes in proteins due to single amino acid replacements, Protein Engineering, Design and Selection 23, 683–387 (2010).

[28] Saraboji K, Gromiha MM, Ponnuswamy MN, Average Assignment Method for Predicting the Stability of Protein Mutants, Biopolymers, 82, 80–92.

[29] Becktel WJ, Schellman JA, Protein Stability Curve, Biopolymers 8, 1859 (1987).

[30] Folch, B, Rooman, M, Dehouck, Y, Modelling Thermal Stability Changes Upon Mutations in Proteins with Artificial Neural Networks, IFAC Proceedings of the 11th International Symposium on Computer Applications in Biotechnology, 11, 525–530 (2010).

[31] Folch B, Rooman M, Dehouck Y, Thermostabilityofsalt bridges versus hydrophobic interactions in proteins probed by statistical potentials, J. Chem. Inf. Model. 48, 119–127 (2008).

[32] Pucci F, Dhanani M, Dehouck Y, Rooman M, Protein Thermostability Prediction within Homologous Families using temperature-dependent statistical potentials, PLoS ONE 9(3), e91659 (2014).

[33] Pucci F, Rooman M, Protein stability curve prediction using temperature-dependent statistical potential, PLoS Comput Biol 10(7), e1003689 (2014).

[34] Tanaka S, Scheraga HA, Medium and long-range interaction parameters between amino acids for predicting three-dimensional structures of proteins, Macromolecules 9, 945–950 (1976).

[35] Miyazawa S., Jernigan R.L., Estimation of effective interresidue contact energies from protein crystal structures: quasi-chemical approximation, Macromolecules 18, 534–552 (1985).

[36] Sippl MJ, Calculation of conformational ensembles from potentials of mean force. An approach to the knowledge based prediction of local structures in globular proteins, J. Mol. Biol. 213, 859–883 (1990).

[37] Thomas PD, Dill KA, Statistical potentials extracted from protein structures: how accurate are they?, J. Mol. Biol. 257,457–69 (1996).

[38] Finkelstein AV, Badretdinov AY, Gutin AM, Why do protein architectures have Boltzmann-like statistics?, Proteins 23, 142–50 (1995).

[39] Li X, Liang J, Knowledge-Based Energy Functions for Computational Studies of Proteins, Computational Methods for Protein Structure Prediction and Modeling, 71–123 (Springer, 2007).

[40] Rooman M, Gilis D, Different derivations of knowledge-based potentials and analysis of their robustness and context-dependent predictive power, Eur. J. Biochem, 254, 135–143 (1998).

[41] Dehouck Y, Gilis D, Rooman M, A new generation of statistical potentials for proteins, Biophys. J. 90, 4010–4017 (2006).

[42] Kocher JP, Rooman M, Wodak S, Factors influencing the ability of knowledge based potentials to identify native sequence-structure matches J. Mol. Biol. 235, 1598–1613 (1994).

[43] Iyer MS, Rhinehart RR, A Method to Determine the Required Number of Neural-Network Training Repetitions, IEEE Transactions on Neural Networks 10, 427–432 (1999).

[44] Atakulreka A, Sutivong D, Avoiding Local Minima in Feedforward Neural Networks by Simultaneous Learning, AI 2007: Advances in Artificial Intelligence, Lecture Notes in Computer Science 4830, 100–109 (2007).

[45] Prechelt L, Neural Networks: Tricks of the trade, 55–69, (Springer Berlin Heidelberg, 1996).

[46] Prechelt L, Automatic early stopping using cross validation: quantifying the criteria. Neural Network 11, 761–767 (1998).

[47] Gilis D, Rooman M, Predicting protein stability changes upon mutation using database-derived potentials: solvent accessibility determines the importance of local versus non-local interactions along the sequence, J. Mol. Biol. 272, 276–290 (1997).

[48] Robertson AD, Murphy KP, Protein Structure and the Energetic of Protein Stability, Chem Rev 97, 1251–1268 (1997).

[49] Bourgeas R, Pucci F, Rooman M, HoTMuSiC v1.0: A webserver for the rational design of proteins with modified thermal resistance, in preparation.

[50] Di Nardo AA, Larson SM, Davidson AR, The Relationship Between Conservation, Thermodynamic Stability, and Function in the SH3 Domain Hydrophobic Core, J. Mol. Biol., 333, 641–655 (2003).

[51] Ratnaparkhi GS, Varadarajan R, Thermodynamic and structural studies of cavity formation in proteins suggest that loss of packing interactions rather than the hydrophobic effect dominates the observed energetics, Biochemistry 39, 12365–12374 (2000).

[52] Eriksson AE et al., Response of a protein structure to cavity-creating mutations and its relation to the hydrophobic effect, Science, 255, 178–183 (1992).

[53] Main ER, Fulton KF, Jackson SE, Context-dependent nature of destabilizing mutations on the stability of FKBP12, Biochemistry 37, 6145–6153 (1998).

[54] Cota E, Hamill SJ, Fowler SB, Clarke J, Two proteins with the same structure respond very differently to mutation: the role of plasticity in protein stability, J. Mol. Biol., 302, 713–725 (2000).

[55] Zavodszky P, Kardos J, Svingor A and Petsko GA, Adjustement of conformational flexibility is a key event in the thermal adaptation of proteins, Proc. Natl. Acad. Sci. USA 95, 7406–7411 (1998).

[56] Kalimeri M, Rahaman O, Melchionna S, Sterpone S, How Conformational Flexibility Stabilizes the Hyperthermophilic Elongation Factor G-domain, J Phys Chem B. 117, 13775–13785 (2013).

[57] Radestock S, Gohlke H, Protein rigidity and thermophilic adaptation, Proteins 79, 1089–108 (2011).

[58] Stafford KA, Robustelli P, Palmer AG, Thermal adaptation of conformational dynamics in ribonuclease H, PLoS Comput Biol. 9, e1003218 (2013).

[59] Kumar MD et al., ProTherm and ProNIT: thermodynamic databases for proteins and protein-nucleic acid interactions, Nucleic Acids Research 34, D204 (2006).

[60] Pucci F, Bourgeas R, Rooman M, High-quality thermodynamic data on the thermal stability changes of proteins upon single-site mutations, Journal of Physical and Chemical Reference Data, submitted (2016), http://dx.doi.org/10.1101/036301.

